# Field Survey of *Anopheles* Mosquito and Their Culturable Gut Bacteria in the Border Regions of Yunnan Province, China

**DOI:** 10.1101/2025.04.02.645363

**Authors:** Guanzhen Fan, Wenxu Yang, Chunli Ding, Peng Tian, Jingwen Wang, Feng Liu, Yaming Yang

**Affiliations:** School of Public Health, Kunming Medical University, Kunming, Yunnan 650500, China; Yunnan Institute of Parasitic Diseases, Pu’er, Yunnan 665000, China; School of Life Sciences, Fudan University, Shanghai 200433, China; Institute of Infectious Diseases, Shenzhen Bay Laboratory, Shenzhen 518132, China

**Keywords:** Yunnan Province, *Anopheles*, Symbiotic Bacteria, Isolation and Identification

## Abstract

Despite the elimination of domestic malaria transmission within China, the border areas of Yunnan Province continue to face a heightened risk of imported cases due to the presence of diverse *Anopheles* mosquito species and the widespread burden of malaria in neighboring countries. While the mosquito gut microbiota offers a promising avenue for developing anti-malaria strategies, a meticulous investigation of the various *Anopheles* species and their associated gut microbiota in Yunnan’s border areas has yet to be undertaken. To address this gap, field surveys were conducted over the years 2020, 2021, and 2023. These surveys resulted in the collection of a substantial sample comprising 4,221 adult female mosquitoes spanning sixteen species with *Anopheles sinensis* identified as the predominant species. Furthermore, 741 cultivable strains of gut symbiotic bacteria were isolated and classified into four phyla comprising 109 genera with the composition of dominant bacteria varying annually. This study provides valuable insights into the population dynamics of malaria vectors in the border regions of Yunnan province, China, along with the intestinal bacteria they harbor including several bacterial species which exhibit promising anti-*plasmodium* effects. By leveraging the antagonistic effects of symbiotic bacteria against *Plasmodium*, our findings contribute to an ongoing global endeavor to develop innovative *Anopheles* management and malaria prevention strategies.

## Introduction

Malaria is a life-threatening disease caused by *Plasmodium* parasites, that are transmitted to humans through the bites of infected *Anopheles* mosquitoes. China successfully implemented a national malaria elimination strategy and achieved complete elimination in 2021 (Cao et al., 2021), which was certified by the World Health Organization (WHO., 2021). However, epidemiological challenges persist in border regions, most notably in Yunnan Province which shares borders with malaria-endemic countries including Myanmar, Laos, and Vietnam (Zhang 2021, Wu 2022, Lin and Zhou 2023). Although local malaria cases have been absent in Yunnan Province since 2017 (Duan et al. 2022), due to the resurgence of malaria in the neighboring regions, 665 imported malaria cases have been reported from 2021 to 2023 (Zhou et al. 2024). In addition, the region’s unique geographical features including low-latitude mountainous terrain (mean annual temperature: 14-33°C), hyper-humid conditions, and dense vegetation provide an ideal ecological environment with optimal microhabitats for *Anopheles* mosquito breeding and proliferation (Atoni et al. 2020).

The vectorial capacity of *Anopheles* mosquitoes is modulated by multiple factors, such as host preference, vector longevity, and gut microbial composition (Narula et al. 2019). Notably, endosymbionts residing in the midgut demonstrate dual regulatory functions. Specific bacterial taxa exhibit either *Plasmodium-*inhibitory or *Plasmodium-*enhancing activities through molecular modulate of the parasite’s developmental cycle (Saraiva 2018, Bai 2019). For instance, several culturable symbiotic bacterial strains—such as *Enterobacter, Serratia ureilytica*, and *Serratia marcescens*—isolated from feild *Anopheles* mosquito populations have demonstrated potent transmission-blocking efficacy (Cirimotich 2011, Gao 2021, 2023).

In contrast, another *Anopheles* mosquito gut bacteria *Asaia bogorensis* helps remodel glucose metabolism in a way that increases midgut pH, thereby promoting *Plasmodium* gametogenesis (Wang et al. 2021). Given the demonstrated potential of targeting symbiotic microbial communities in mosquito midguts for malaria control, comprehensive field surveys of gut microbiota composition across species are critically important for the development of effective anti-malaria strategies. Moreover, these detailed studies are especially important considering the gut microbiota can be highly variable between and within mosquito species due to diverse ecological factors such as nutrient availability, microclimatic conditions, and blood meal sources (Coon 2016, Gimonneau 2014).

Yunnan Province was historically a high-endemic area for malaria, posing a significant public health threat (Duan et al. 2022). Despite China’s achievement in domestic malaria elimination, hyperendemic persistence in bordering nations (Myanmar, Vietnam, and Laos) sustains significant epidemiological pressure. The *Anopheles* vector complex remains a persistent regional threat due to its transnational transmission potential. As a result, Yunnan Province harbors the most diverse assemblage of *Anopheles* species in China. However, a thorough investigation of *Anopheles* mosquito species and their gut microbiota in the border area of Yunnan Province is lacking. To address this gap, field surveys were conducted during peak mosquito density seasons (July–August) in 2020, 2021, and 2023 across three border counties: Yingjiang (China–Myanmar border), Mengla (China–Laos border), and Jinping (China–Vietnam border). In parallel with *Anopheles* species investigations, culturable gut symbiotic bacteria were isolated and identified.

## Methods

### Field Collection of *Anopheles* Mosquitoes

Based on China’s border regions adjacent to the Mekong River Basin, combined with confidential data on historical *Anopheles* mosquito distribution, we selected three county-level survey locations: Yingjiang County (Dehong Dai and Jingpo Autonomous Prefecture, bordering Myanmar); Mengla County (Xishuangbanna Dai Autonomous Prefecture, bordering Laos); and Jinping County (Honghe Hani and Yi Autonomous Prefecture, bordering Vietnam) (Yang 2021). These selected natural village survey locations all met specific criteria for *Anopheles* mosquito breeding, such as the presence of cattle sheds and nearby rice paddies. A total of 10 survey sites were utilized across these three counties (**Figure 1**, the detailed information of these locations was listed in **Supp. Table S1**). Field collections were carried out during the peak *Anopheles* mosquito activity season (July–August) in 2020, 2021, and 2023 using two collection methods: an aspirator-based method for capturing mosquitoes from cattle sheds (19:00–24:00) (Yin et al. 1993) and a mosquito trap lamp placed overnight from 19:00 to 7:00 (Zhou et al. 2001). Distinguish between female and male mosquitoes based on their external characteristics such as their size and antennae All of the collected *Anopheles* adult female mosquitoes were identified morphologically to the species level using a taxomic identification key (The Identification Key to Chinese *Anopheles*).

**Figure 1.**
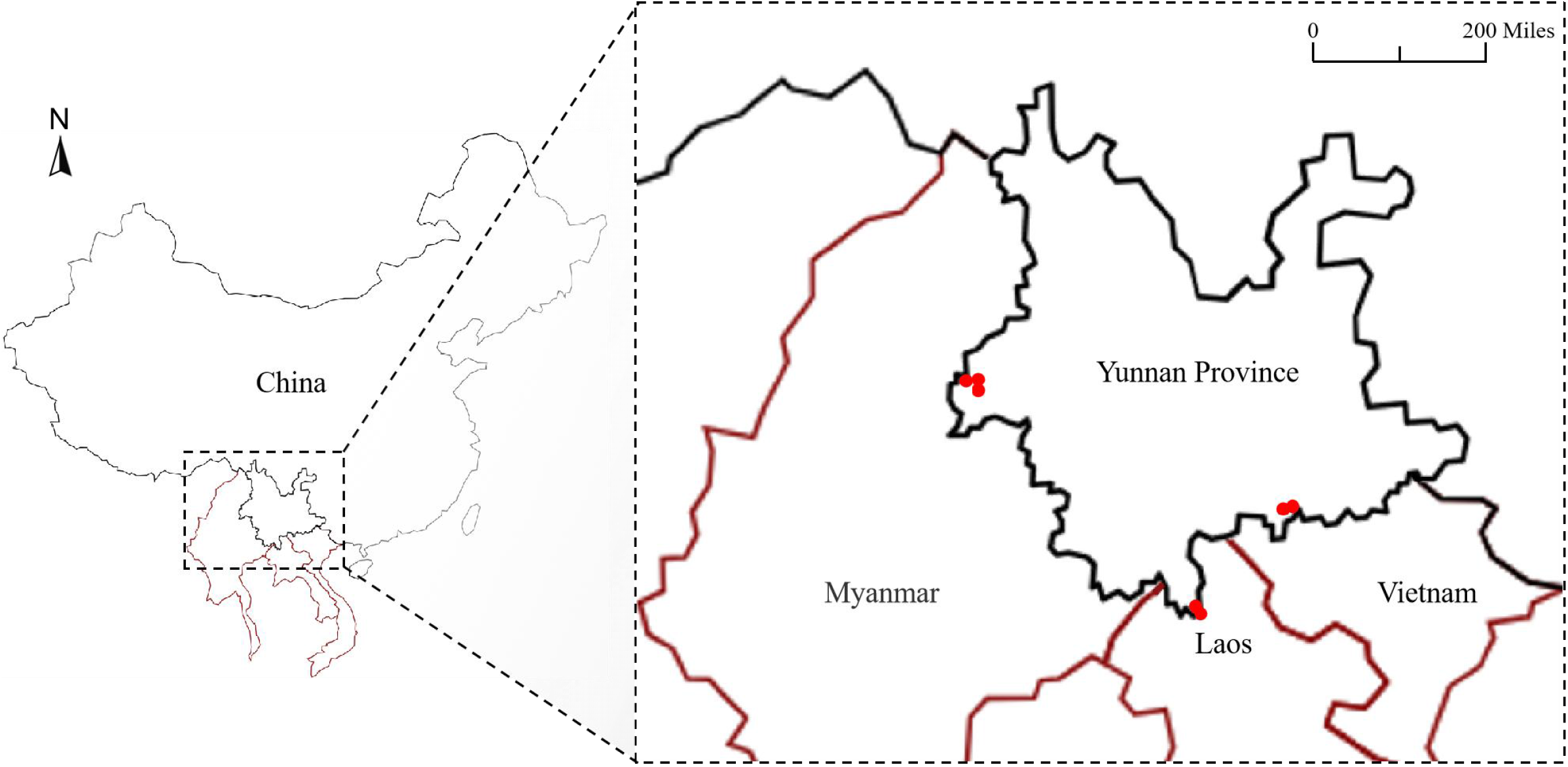
Survey locations adjacent to border countries in Yunnan Province (2020-2021 and 2023). The red dots represent the coordinates of the sampling location.

### Bacterial isolates and routine culture

Specimens preparation and dissection were performed following previously described methods (Chandel et al. 2013). Briefly, the isolated midguts were homogenized in normal saline and serially diluted to 0.1% (v/v). The diluted homogenates were plated on different culture media: Luria-Bertani agar medium, Blood agar medium, or MRS agar medium. Plates were then incubated at 28°C for 48 hours. Based on their growth characteristics (i.e., colony morphology, size, color, and growth margins), individual colonies were selected and subjected to streaking for further subculture to obtain monoclonal isolations.

### 16S rRNA Gene Amplification and Sequencing

The full-length 16S rRNA gene (∼1500 bp) was amplified using universal primers: 27F (5’-AGAGTTTGATCCTGGCTCAG-3’) and 1492R (5’-GGTTACCTTGTTACGACTT-3’). The cycling conditions included an initial denaturation at 94°C for 5 minutes, followed by 35 cycles of denaturation at 94°C for 30 seconds, annealing at 57°C for 30 seconds, and extension at 72°C for 1.5 minute, and final extension at 72°C for 10 minutes. The amplified target fragments were purified and ligated into pMD19-T vector and then transformed into *Escherichia coli* DH5α competent cells and sent for sequencing. The obtained sequences were compared with the NCBI database using BLASTn to identify the most similar bacteria sequences.

## Results

### Population structure of captured *Anopheles* mosquito species

*Anopheles* mosquitoes were collected from 10 survey sites across Yingjiang County, Mengla County, and Jinping County in Yunnan Province during 2020, 2021, and 2023. Field collections were not performed in the year 2022due to COVID-19 travel restrictions in China. A total of 505, 1995, and 1721 *Anopheles* mosquitoes, representing 16 different species, were captured in 2020, 2021 and 2023, respectively (**Table 1**). The identified species included *Anopheles sinensis, Anopheles minimus, Anopheles vagus, Anopheles peditaeniatus, Anopheles kunmingensis, Anopheles tessellatus, Anopheles argyropus, Anopheles pseudowillmori, Anopheles crawfordi, Anopheles splendidus, Anopheles maculatus, Anopheles jeyporiensis, Anopheles culicifacies, Anopheles barbirostris, Anopheles kochi, Anopheles acunitus*. Among the collected specimens, *An. sinensis* represented the most abundant species with the largest average sampling ratio of 50.86%. Other prevalent species included *An. kochi, An. vagus, An. peditaeniatus, An. kunmingensis, An. minimus*, and *An. tessellatus*, with moderate abundances ranging from 5.95% to 9.55% in our total collection. All of the other nine *Anopheles* mosquito species have low prevalence (capturing ratio <1%) in our sampling location and were considered to be minor species in the border region of Yunnan Province (**Table 2**).

**Table 1.** The total number of *Anopheles* mosquitoes captured within the border counties of Yunnan Province (2020-2021 and 2023). The numbers in the table represent the number of captured mosquitoes, and “-” indicates that no mosquitoes of that species were captured.

| Mosquito species | Yingjiang County |  |  | Mengla County |  |  | Jinping County |  |  |
| --- | --- | --- | --- | --- | --- | --- | --- | --- | --- |
|  | 2020 | 2021 | 2023 | 2020 | 2021 | 2023 | 2020 | 2021 | 2023 |
| *An. sinensis* | 51 | 132 | 443 | 66 | 393 | 246 | 51 | 259 | 506 |
| *An. minimus* | 53 | 3 | 1 | - | - | 2 | 35 | 130 | 27 |
| *An. peditaeniatus* | 20 | 208 | 1 | 20 | 1 | - | 5 | 1 | - |
| *An. pseudowillmori* | - | - | - | - | - | - | 7 | 13 | - |
| *An. vagus* | - | - | - | 71 | 140 | 46 | 8 | 73 | 1 |
| *An. jeyporiensis* | - | - | - | - | - | - | 1 | - | 23 |
| *An. argyropus* | - | - | - | 6 | 8 | - | - | 12 | - |
| *An. tessellatus* | - | - | - | 42 | 103 | 104 | - | 4 | 10 |
| *An. kochi* | - | - | 1 | 21 | 307 | 66 | - | 2 | 6 |
| *An. maculatus* | - | - | - | - | - | - | - | - | 73 |
| *An. crawfordi* | - | - | - | 6 | 4 | - | - | - | - |
| *An. barbirostris* | - | - | - | 2 | 1 | 6 | - | - | - |

| Mosquito<br>species | Yingjiang County |  |  | Mengla County |  |  | Jinping County |  |  |
| --- | --- | --- | --- | --- | --- | --- | --- | --- | --- |
|  | 2020 | 2021 | 2023 | 2020 | 2021 | 2023 | 2020 | 2021 | 2023 |
| *An. jeyporiensis* | - | - | - | - | - | 23 | - | - | - |
| *An. aconitus* | - | - | - | - | - | 2 | - | - | - |
| *An. splendidus* | 2 | - | - | - | - | - | - | - | - |
| *An. kunmingensis* | 37 | 201 | 134 | - | - | - | - | - | - |
| *An. culicifacies* | 1 | - | - | - | - | - | - | - | - |
| Total | 164 | 544 | 580 | 234 | 957 | 495 | 107 | 494 | 646 |

**Table 2.**
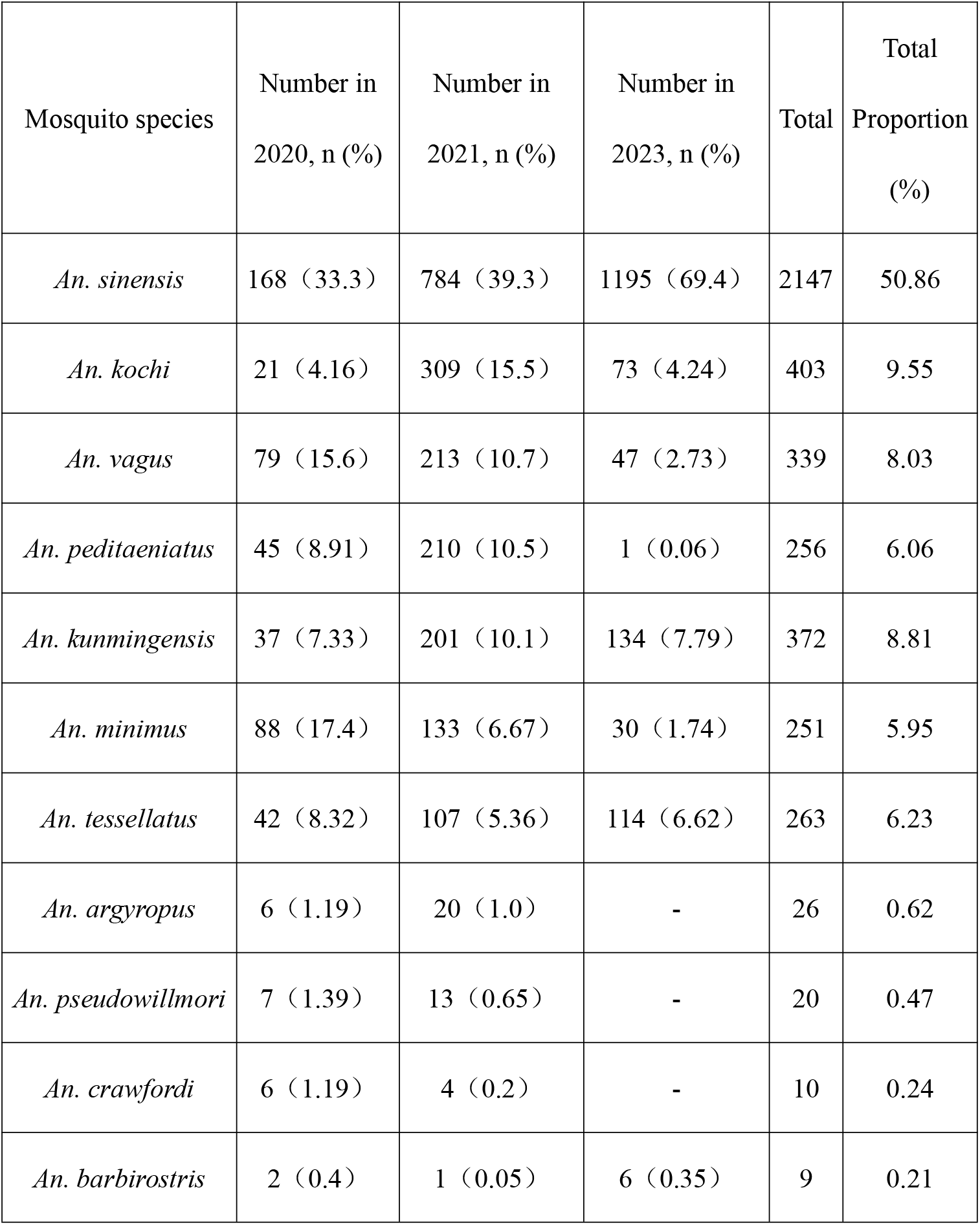
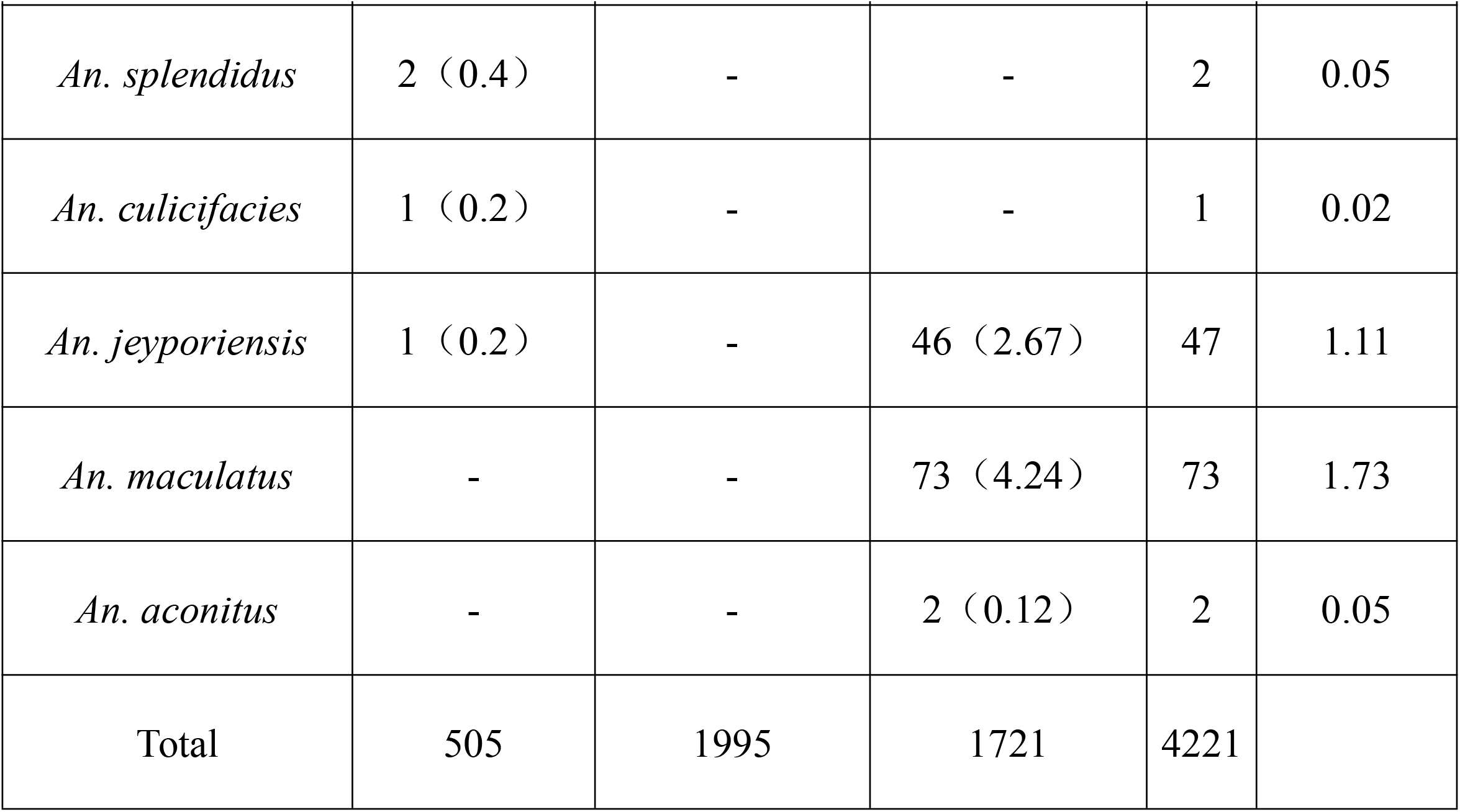
Composition of *Anopheles* mosquitoes within the border counties of Yunnan Province (2020-2021 and 2023). Numbers outside the parentheses represent the number of captured mosquitoes, and the number inside the parentheses indicates the proportion of mosquitoes. “-” indicates that no mosquitoes of that species were captured.

The population structure of *Anopheles* species exhibited substantial variation across different sampling years. For example, while *An. sinensis* remained the most predominant species throughout all three years, its relative abundance increased significantly in 2023, accounting for nearly 70% of the total collection, compared to 33.3% in 2020 and 39.3% in 2021. In contrast, *An. peditaeniatus* and *An. minimus*, which were moderately abundant in 2020 and 2021, were found in markedly lower numbers in 2023. Moreover, we also noticed that some low prevalence mosquito species—such as *An. argyropus, An. pseudowillmori* and *An. Crawfordi*—were identified in 2020 and 2021 but were absent in 2023 even though the total number of mosquitoes collected in 2023 was comparable with that in 2021.

### Identification of culturable midgut symbiotic bacteria in wild *Anopheles* mosquitoes

Following taxonomic identification, wild-caught mosquitoes were sacrificed for midgut symbiotic bacteria isolation. The isolated bacteria were transferred to growing plates with different culture media under appropriate temperature. After successful laboratory culturing, the bacteria were identified through 16S rRNA gene sequencing. A total of 741 bacterial strains were identified with 234 strains in 2020, 293 strains in 2021, and 214 strains in 2023 which distributed in four phyla, including Proteobacteria, Actinobacteria, Bacteroidetes, and Firmicutes (**Table 3**). Among them, Proteobacteria was the predominant phylum of the culturable bacteria isolated from all mosquitoes comprising more than 60% of the strains identified in our samples. The second most abundant bacteria specie was classified as Actinobacteria, which account for 18.89% of the total sampling. In contrast, Bacteroidetes or Firmicutes were less abundant, comprising around 10% of the culturable bacterial strains.

**Table 3:** Phylum-level identification of culturable gut symbiotic bacteria in wild *Anopheles* mosquitoes within the border counties of Yunnan Province (2020-2021 and 2023). The data represent the number of bacterial strains at the phylum level in the gut microbiota of *Anopheles* mosquitoes.

| Phylum Level | 2020 | 2021 | 2023 |
| --- | --- | --- | --- |
| Proteobacteria | 167 | 146 | 138 |
| Actinobacteria | 34 | 82 | 24 |
| Bacteroidetes | 21 | 13 | 29 |
| Firmicutes | 12 | 52 | 23 |
| total | 234 | 293 | 214 |

To get more detailed information about the symbiotic bacterial communities in the mosquito midgut, we further classified the 741 culturable strains into 109 genera. Among them, 23 genera with a quantity greater than or equal to 5 have been identified, including *Thorsellia, Brevundimonas, Cupriavidus, Herbaspirillum, Enterobacter, Pantoea, Serratia, Asaia, Agrobacterium, Stenotrophomonas, Pseudomonas, Delftia, Acinetobacter, Sphingomonas, Microbacterium, Curtobacterium, Bacillus, Micrococcus, Staphylococcus, Enterococcus, Paenibacillus, Sphingobacterium*, and *Chryseobacterium*. Notably, *Moraxellaceae acinetobacter* and *Microbacteriaceae* m*icrobacterium* were the most common cultivable commensal bacteria in the midgut from the Yunnan border area as surveyed, with a total of 89 and 76 strains, accounting for 12.01% and 10% of all isolates, respectively (**Table 4**). However, the predominant culturable symbiotic bacteria varied across sampling years, indicating dynamic microbiome compositions. For example, in 2020, *M. acinetobacter* was the predominant genus, accounting for 24.36% of the isolated strains while in 2021, *M. microbacterium* was the most abundant, with a proportion of 14.68%. In 2023, *Thorselliaceae thorsellia* (17.29%) and *Weeksellaceae chryseobacterium* (11.68%) became the predominant genera (**Table 5**). These findings highlight the temporal variability and dynamic nature of the midgut microbiome in *Anopheles* mosquitoes of the Yunnan border areas.

**Table 4:** Genus-level identification of culturable gut symbiotic bacteria in wild *Anopheles* mosquitoes within the border counties of Yunnan Province (2020-2021 and 2023). Others: the total number of identified symbiotic bacterial genera is less than 5.

| Phylum | Family | Geneus | Number | Total Proportion (%) |
| --- | --- | --- | --- | --- |
| Proteobacteria | Thorselliaceae | *Thorsellia* | 53 | 7.15 |
|  | Caulobacteraceae | *Brevundimonas* | 25 | 3.37 |
|  | Burkholderiaceae | *Cupriavidus* | 7 | 0.94 |
|  | Oxalobacteraceae | *Herbaspirillum* | 10 | 1.35 |
|  | Enterobacteriaceae | *Enterobacter* | 10 | 1.35 |
|  | Erwiniaceae | *Pantoea* | 10 | 1.35 |
|  | Yersiniaceae | *Serratia* | 15 | 2.02 |
|  | Acetobacteraceae | *Asaia* | 9 | 1.21 |
|  | Rhizobiaceae | *Agrobacterium* | 34 | 4.59 |
|  | Lysobacteraceae | *Stenotrophomonas* | 20 | 2.70 |
|  | Pseudomonadaceae | *Pseudomonas* | 43 | 5.80 |
|  | Comamonadaceae | *Delftia* | 6 | 0.81 |
|  | Moraxellaceae | *Acinetobacter* | 89 | 12.01 |
|  | Sphingomonadaceae | *Sphingomonas* | 13 | 1.75 |
| Actinobacteria | Microbacteriaceae | *Microbacterium* | 76 | 10.26 |
| Actinobacteria | Microbacteriaceae | *Curtobacterium* | 7 | 0.94 |
| Firmicutes | Bacillaceae | *Bacillus* | 32 | 4.32 |
|  | Micrococcaceae | *Micrococcus* | 5 | 0.68 |
|  | Staphylococcaceae | *Staphylococcus* | 18 | 2.43 |
|  | Enterococcaceae | *Enterococcus* | 7 | 0.94 |
|  | Paenibacillaceae | *Paenibacillus* | 6 | 0.81 |
| Bacteroidetes | Sphingobacteriaceae | *Sphingobacterium* | 14 | 1.89 |
|  | Weeksellaceae | *Chryseobacterium* | 37 | 4.99 |
| others |  | others | 195 | 26.32 |
|  |  | Total | 741 |  |

**Table 5:** Proportion of gut symbiotic bacteria in wild *Anopheles* mosquitoes from the border counties of Yunnan Province. Others: the total number of identified symbiotic bacterial genera is less than 5.

| Survey Year | Geneus | Number | Total Proportion (%) |
| --- | --- | --- | --- |
| 2020 | *Acinetobacter* | 57 | 24.36 |
|  | *Pseudomonas* | 23 | 9.83 |
|  | *Brevundimonas* | 19 | 8.12 |
|  | *Microbacterium* | 17 | 7.26 |
|  | *Sphingobacterium* | 14 | 5.98 |
|  | *Sphingomonas* | 13 | 5.56 |
|  | *Stenotrophomonas* | 8 | 3.42 |
|  | *Thorsellia* | 7 | 2.99 |
|  | *Chryseobacterium* | 7 | 2.99 |
|  | *Cupriavidus* | 7 | 2.99 |
|  | *Bacillus* | 7 | 2.99 |
|  | *Delftia* | 6 | 2.56 |
|  | *Micrococcus* | 5 | 2.14 |
|  | Others | 44 | 18.80 |
| 2021 | *Microbacterium* | 43 | 14.68 |
|  | *Agrobacterium* | 27 | 9.22 |
|  | *Acinetobacter* | 26 | 8.87 |
|  | *Staphylococcus* | 18 | 6.14 |
| 2021 | *Bacillus* | 12 | 4.10 |
|  | *Herbaspirillum* | 10 | 3.41 |
|  | *Pantoea* | 10 | 3.41 |
|  | *Pseudomonas* | 10 | 3.41 |
|  | *Serratia* | 10 | 3.41 |
|  | *Thorsellia* | 9 | 3.07 |
|  | *Enterococcus* | 7 | 2.39 |
|  | *Curtobacterium* | 7 | 2.39 |
|  | *Brevundimonas* | 6 | 2.05 |
|  | *Paenibacillus* | 6 | 2.05 |
|  | *Chryseobacterium* | 5 | 1.71 |
|  | Others | 87 | 29.69 |
| 2023 | *Thorsellia* | 37 | 17.29 |
|  | *Chryseobacterium* | 25 | 11.68 |
|  | *Microbacterium* | 16 | 7.48 |
|  | *Bacillus* | 13 | 6.07 |
|  | *Stenotrophomonas* | 12 | 5.61 |
|  | *Enterobacter* | 10 | 4.67 |
|  | *Pseudomonas* | 10 | 4.67 |
|  | *Asaia* | 9 | 4.21 |
|  | *Agrobacterium* | 7 | 3.27 |
| 2023 | *Acinetobacter* | 6 | 2.80 |
|  | *Serratia* | 5 | 2.34 |
|  | Others | 64 | 29.91 |

## Discussion

As reported in previous studies, the four primary malaria vectors in China are *An. sinensis, An. minimus, An. dirus*, and *An. lesteri*, while the significant malaria vectors in Yunnan Province include *An. sinensis, An. minimus*, and *An. jeyporiensis* (Dong 2000). Consistent with these findings, our survey indicates that *An. sinensis* is the most abundant *Anopheles* species in the Yunnan border region, aligning with observations at both the national and provincial levels (**Table 2**). However, we also notice some unique features of the population structures of *Anopheles* mosquitoes. While *An. minimus* and *An. jeyporiensis* were reportedly the most populous malaria vector in Yunnan province, these species only comprise a small percentage (6% and 1%, respectively) of our sampled mosquitoes, suggesting the composition of mosquito species may vary in the border regions compared to the general constitution of *Anopheles* species in the entire province. This may also simply be due to the collection methods. Different species may be easier or harder to capture depending on the method used, time of day, time of year, temperature (Kelly-Hope 2009, Ndiath 2011). Moreover, *An. dirus* and *An. lesteri*, which represent the third and fourth most dominant malaria vectors in China, respectively, were not present in our survey at all, which may be due to their particular requirement for living habitats such as vegetation coverage, temperature, and humidity. However, these discrepancies may also stem from differences in collection methods, as species-specific capture efficiency can be influenced by factors such as trapping technique, time of day, seasonality, and environmental conditions like temperature and humidity.

Over evolutionary time, *Anopheles* mosquitoes have established intricate symbiotic relationships with their commensal microbiota and the *Plasmodium* parasite. The mosquito’s midgut serves as the initial site for *Plasmodium* transmission, where the local micro-environment and gut microbiota play critical roles in parasite survival, development, and subsequent maturation (Bando et al. 2013). Previous studies demonstrated specific gut-associated commensal bacteria isolated from *Anopheles* mosquitoes have the potential to influence *Plasmodium* transmission. For example, *Pseudomonas* species isolated from *Anopheles* mosquitoes in Shanghai, China, have been shown to catabolize 3-hydroxykynurenine (3-HK), thereby suppressing *Plasmodium* development in the mosquito midgut and reducing malaria transmission (Feng et al. 2022). Interestingly, *Pseudomonas* species were also identified among the culturable strains in our survey, suggesting their stable constitutional colonization in different *Anopheles* mosquito. It is tempting for us to speculate that there may be additional bacteria strains in the total collections we isolated in this study that also exert anti-*plasmodium* effects which would represent promising targets for malaria control.

In this study, a moderate number of commensal bacterial strains were isolated, with several bacterial genera identified in *Anopheles* mosquitoes also being commonly found in other mosquito species. Recent studies have reported the presence of *Microbacterium* across all developmental stages of *Aedes albopictus* (Zhao et al. 2022). Similarly, *Microbacterium* has been isolated from laboratory-reared strains of *Culex pipiens pallens* (Lyu et al. 2023). Another frequently detected symbiotic bacterium in the midgut of *Anopheles* mosquitoes is *Thorsellia*, which has been isolated from other *Anopheles* species such as *Anopheles. gambiae* (Dillon 1991, Briones 2008), *Anopheles. stephensi* (Rani et al. 2009), *An. culicifacies* (Chavshin et al. 2014), and *Anopheles. arabiensis* (Kampfer et al. 2015). Notably, *Thorsellia* has been detected not only in the midgut of *An. gambiae* but also in its larval aquatic habitat (Briones et al. 2008), suggesting that adult *Anopheles* mosquitoes may acquire these bacteria during the larva stage from *Thorsellia*-enriched aquatic environment. Future investigation of microbial organisms in the aquatic habitats of mosquito larvae may give us more information about the origins and transmission pathways of commensal bacteria in adult *Anopheles* mosquitoes.

In addition, traditional culture-dependent methods for isolating commensal bacteria present inherent technical limitations, particularly in the cultivation of anaerobic bacteria, making it difficult to fully cultivate all gut cultivable commensal microbiota. While the bacterial profiles obtained in this study may not fully represent the diversity of midgut commensal bacteria in wild mosquitoes, these findings provide a preliminary understanding of the culturable gut microbiota composition in *Anopheles* mosquitoes from the border regions of Yunnan Province, offering insights for future exploitation and potential applications that target gut commensal bacteria. Future studies should aim to explore the diversity of commensal bacteria in the guts of *Anopheles* mosquitoes with more sophisticated technologies, such as combining high-throughput sequencing-dependent methods (like 16S amplicon analysis) and metagenomics analysis, which should provide a more complete picture of the commensal bacteria of wild *Anopheles* mosquitoes in the border regions of Yunnan Province, China.

## Supporting information

Supplemental Table1, and will be used for the link to the file on the preprint site.

## Author contributions

Guanzhen Fan: Conceptualization, Methodology, Data Curation, Investigation, Formal

Analysis, Writing - Original Draft.

Wenxu Yang: Investigation, Data Curation, Writing - Original Draft.

Chunli Ding: Visualization, Investigation.

Peng Tian: Visualization, Investigation.

Jingwen Wang: Conceptualization, Supervision, Resources.

Feng Liu: Resource, Validation, Writing - Review & Editing.

Yaming Yang: Conceptualization, Funding Acquisition, Project Administration, Resources, Supervision, Writing - Review & Editing.

## Acknowledgments

We thank all members of the Y.Y. W.J. and L.F. lab for suggestions and comments. We thank Kunming Medical University and Fudan University for equipment support. This work was funded by National Natural Science Foundation of China (Grant No. U1902211).

