## Supplemental Table1, and will be used for the link to the file on the preprint site. for "Field Survey of *Anopheles* Mosquito and Their Culturable Gut Bacteria in the Border Regions of Yunnan Province, China"

Table S1. Overview of sampling sites within the border counties of Yunnan Province (2020-2021 and 2023).

| County | Township | Village | Altitude (meters) | Location | Surrounding Agricultural Crop Types | Survey Year |
|---|---|---|---|---|---|---|
| Yingjiang County | Nabang Town | New National Gate | 220 | 97°33'39"N 24°45'60"E | Banana | 2020 |
| | Pingyuan Town | Gaoli Village | 834 | 97°55'15"N 24°47'13"E | Maize, Sugarcane | 2020, 2021 |
| | Xima Town | Zhuanpozhai Village | 1648 | 97°42'14"N 24°46'29"E | Rice, Maize, Rapeseed | 2020, 2021, 2023 |
| | Nongzhang Town | Nansuan Village | 818 | 97°55'48"N 24°37'46"E | Sugarcane, Rice | 2021 |
| | Tongbiguan Township | Nanling Village | 1324 | 97°41'23"N 24°35'25"E | Rice, Maize | 2023 |
| | Tongbiguan Township | Dazhai Village | 1313 | 97°40'30"N 24°37'50"E | Rice | 2023 |
| Mengla County | Shangyong Town | Shanggang Village | 736 | 101°43'27"N 21°16'25"E | Maize, Rubber Trees | 2021, 2023 |

| County | Township | Village | Altitude (meters) | Location | Surrounding Agricultural Crop Types | Survey Year |
|---|---|---|---|---|---|---|
| Mengla County | Shangyong Town | Mozheng Village | 752 | 101°45'90"N 21°11'46"E | Maize, Rubber Trees, Dragon Fruit | 2020, 2021 |
| Jinping County | Jinhe Town | Dabaozhai Village | 1443 | 103°13'17"N 22°49'40"E | Rice, Banana, Maize | 2020, 2021, 2023 |
| | Tongchang Township | Dongzonghe Village | 1080 | 103°4'51"N 22°45'54"E | Rice, Maize, Lemon Trees | 2020, 2021, 2023 |
